# Investigating the neural basis of schematic false memories by examining schematic and lure pattern similarity

**DOI:** 10.1101/2023.07.26.550683

**Authors:** Catherine M. Carpenter, Nancy A. Dennis

## Abstract

Schemas allow us to make assumptions about the world based upon previous experiences and aid in memory organization and retrieval. However, a reliance on schemas may also result in increased false memories to schematically related lures. Prior neuroimaging work has linked schematic processing in memory tasks to activity in prefrontal, visual, and temporal regions. Yet, it is unclear what type of processing in these regions underlies memory errors. The current study examined where schematic lures exhibit greater neural similarity to schematic targets, leading to this memory error, as compared to neural overlap with non-schematic lures, which, like schematic lures, are novel items at retrieval. Results showed that patterns of neural activity in ventromedial prefrontal cortex, medial frontal gyrus, middle temporal gyrus, hippocampus, and occipital cortices exhibited greater neural pattern similarity for schematic targets and schematic lures than between schematic lures and non-schematic lures. As such, results suggest that schematic membership, and not object history, may be more critical to the neural processes underlying memory retrieval in the context of a strong schema.

## Introduction

Schemas provide us with a means for organizing the world around us and allow us to make inferences in new environments based upon previous experiences. In doing so, schemas aid in how we organize, encode, and retrieve information across a multitude of experiences. While schemas generally provide a large benefit to memory, schemas may also result in increased rates of false memories, particularly when novel information matches the schema (Lampinen et al., 2001). In fact, it is relatively common to find comparable true and false memory rates for schematic information over and above that of novel information (Lampinen et al., 2000; Miller & Gazzaniga, 1998; Neuschatz et al., 2002; Webb et al., 2016). This is interesting given that in visual memory paradigms assessing schematic memory, the schematic targets have been presented and studied whereas the schematic lures are novel objects with no perceptual overlap with the targets. In this sense, schematic lures are similar to non-schematic lures which have much lower false memory rates (Lampinen et al., 2000; Miller & Gazzaniga, 1998; Neuschatz et al., 2002; Webb et al., 2016; Webb & Dennis, 2020). While there has been extensive behavioral and quantitative univariate neural work in the realm of schematic false memory (Brewer & Treyens, 1981; Charlton & Leov, 2021; Kleider et al., 2008; Lampinen et al., 2001; Neuschatz et al., 2002; Webb & Dennis, 2020), the underlying neural mechanisms that promote schema-related increases in false memories to these visually distinct objects are less understood. Given the schematic relationship but visual distinction between schematic targets and lures in a visual memory study, we investigated whether the neural mechanisms underlying schematic lures resemble that of target items that share an overlapping schematic theme or whether they reflect that of novel, unique visual objects.

Behavioral evidence is largely conclusive that when schemas are used to support memory, schema-consistent information is well remembered (Alba & Hasher, 1983; Castel, 2005; Spalding et al., 2015; van Kesteren et al., 2012). Schemas act as a guide for integrating new memories and knowledge with prior experiences to form a holistic representation of an experience (Alba & Hasher, 1983). As such, schemas provide a scaffold and support for memory retrieval (van Kesteren et al., 2012). At the same time, these processes lead to schematic information not presented in the studied environment to also be erroneously endorsed as “old” in subsequent memory tests due to the overlap in the gist of prior experiences (e.g., Brewer & Treyens, 1981; Charlton & Leov, 2021; Kleider et al., 2008; Lampinen et al., 2001; Neuschatz et al., 2002; Webb & Dennis, 2020). For example, in the famous “room schema” study by Lampinen and colleagues (2001), participants exhibit higher false recall for schematically-related items found in an office (e.g., books) compared to atypical office items (e.g., mirror). The occurrence of high false memories rates has since been shown in numerous studies using both recall and recognition testing procedures (Charlton & Leov, 2021; Kleider et al., 2008; Lew & Howe, 2017; Neuschatz et al., 2002). Yet false memory rates amongst schematic images often fall below that of semantically related words (Coane et al., 2021; Dennis et al., 2015; Oliver et al., 2016). This may be related to the unique physical properties of schematic lure objects compared to their studied counterparts. In this study we ask what neural processing leads to lures being erroneously endorsed as “old” opposed to correctly being identified as novel objects during memory retrieval.

The fact that schematic lures are unique objects that, despite their conceptual link to materials studied, share no perceptual overlap with studied objects presents a unique question for how these objects are represented in memory. On one hand, schematic lures are highly related to the studied material. For example, a sink is highly related to the concept of bathroom and is conceptually related to other objects in a bathroom schema such as toilet, shower, bathtub, and plunger. Yet schematic lures are also physically distinct from these objects, sharing no physical characteristics with them. Any false alarms to schematically related lures are therefore likely based upon that conceptual relationship, whereas perceptual features likely help to distinguish the object as a novel item during a memory test. Thus, from a neural perspective, this should create a dissociation in neural representations where schematic lures might resemble targets in regions processing schematic information, but not in regions that process visual details of objects in memory tasks.

Past neuroimaging work examining univariate activity has identified a role of several regions, including the ventromedial prefrontal cortex (vmPFC) and middle temporal gyrus (MTG), in supporting true schematic memories (Gilboa & Marlatte, 2017; van Kesteren et al., 2012, 2013; Webb et al., 2016). With respect to the vmPFC, this work posits that while encoding of novel information is initially medial temporal lobe (MTL) dependent, reactivation of schemas will result in a dependency on prefrontal cortices, with consistent schema-related activation occurring in the vmPFC (Brod et al., 2017; Gilboa & Marlatte, 2017; van Kesteren et al., 2010, 2012, 2013). Thus, we may utilize the vmPFC in addition to MTL regions to consolidate and process schema-consistent information if a prior schema exists (van Kesteren et al., 2012; Zeithamova et al., 2012). Specifically, researchers find that having a prior schema facilitates processing in the vmPFC and that increased coupling between the hippocampus (HC) and vmPFC is related to better memory performance for schematic information (van Kesteren et al., 2010). Interestingly, individuals with damage to the vmPFC exhibit a reduced reliance on schema-based processing (Spalding et al., 2015), as well as a reduction in schema-related false memories (Warren et al., 2014). In addition to the supporting role of the vmPFC in true schematic memories, this work highlights a potential role of the vmPFC in promoting false memories to schema-consistent information. Accordingly, a nearby region, the medial prefrontal cortex (mPFC), which contains the middle frontal gyrus (MFG), has also been implicated in many studies of false memories, including schematic, perceptual, and semantically related false alarms (e.g., Abe et al., 2008; Dennis & Turney, 2018; Garoff-Eaton et al., 2007; for a meta-analysis see Kurkela & Dennis, 2016; Slotnick & Schacter, 2006; Turney & Dennis, 2017; Webb & Dennis, 2019). Taken together, results suggest that the schema-related activity in the mPFC may be a key component to the encoding and endorsement of false memories, particularly when the lure information is schematically or thematically related to studied information.

The middle temporal gyrus (MTG) is another region identified in numerous false memory studies, including those associated with semantic, schematic, and perceptual false memories (e.g., Dennis et al., 2007, 2008; Garoff-Eaton et al., 2006; Gilboa & Marlatte, 2017; Kubota et al., 2006; Slotnick & Schacter, 2006; Turney & Dennis, 2017; Webb et al., 2016; Webb & Dennis, 2019). In terms of studies examining schematic false memory, this region has been suggested to represent semantic gist related to the connection between the schema and the lure items (Webb et al., 2016; Webb & Dennis, 2019). Specifically, prior work finds that activation in the MTG is associated with both true and false schematic memories, including highly confident recollection-related memory errors and increasing in activation with regard to individual differences in false memory rates (Dennis & Turney, 2018; Webb et al., 2016). This work highlights the involvement of the MTG in semantic processing and how it may lead to the erroneous endorsement of lures based upon evaluating congruent information that is consistent with prior schemas. The above regions are clearly active when one makes an erroneous “old” endorsement to a lure, however, it is unclear whether the underlying neural processing truly reflects that of target items.

While the foregoing regions have been found to process schematic properties of objects in a memory task, the visual cortex is known for its ability to discriminate between old and new information (Bowman & Dennis, 2015b; Kurkela & Dennis, 2016; Slotnick & Schacter, 2006). Specifically, the visual cortices exhibit unique patterns of neural activation during both perceptual processing (Haxby et al., 2001) and memory retrieval (Bowman & Dennis, 2015a, 2015b; Slotnick & Schacter, 2004, 2006; Stark et al., 2010; Vaidya et al., 2002; Webb et al., 2016; Wheeler et al., 2000) when objects are physically distinct from one another. Regarding false memory, while the visual cortex has shown greater similarity in processing perceptually related information (Bowman & Dennis, 2015a; Gutchess & Schacter, 2012), it has also been implicated in the correct rejection of lures that are more visually distinct from that presented during encoding (Bowman & Dennis, 2015a, 2015b). This suggests that the visual cortex is sensitive to different types of novelty processing and may support successful memory discrimination for semantically related, but perceptually novel, information (Bowman & Dennis, 2015a). Schematically related information should fall in this domain, being physically and perceptually unique in its composition compared to related items from the encoded schema. With regards to the visual cortex’s role in both successful retrieval of schematic information and correct rejection of novel and related items, it is worthwhile to consider the underlying quality of relationships in visual regions with regards to how they process schematically related and novel information, and how that may relate to memory success or failure.

The current work aims to elucidate how schematic lures are represented neurally, specifically examining the similarity of neural patterns with that of both schematic target and novel lures. While prior work has developed a greater understanding of the quantitative relationships of blood-oxygen-level-dependent (BOLD) activation associated with schematic processing, the qualitative relationships underlying those patterns has remained an open question. The purpose of our investigation is to examine the quality of these relationships using pattern similarity analyses to determine whether—and where, neurally—schematic lures are represented more similar to that of studied, schematic information or more similar to other unstudied novel information. In addition to elucidating the neural representations of schematic information, we are interested in examining whether these patterns correlate with false alarm rates. In the context of the current set of analyses we define schema as a cognitive framework that help us to organize, interpret, and synthesize information. With respect to the current design, we further operationalize a schema as a scene depicting a singular concept such as ‘bathroom’ or ‘farm’. Included in the scene are objects that are best characterized by their membership in the schema. For example, a toilet is (typically) found in no other schema but that of bathroom (see Methods and Appendix A for additional details on schema inclusion).

We hypothesize that schematic information will have more similar neural patterns to one another irrespective of whether it was a target or lure trial at retrieval (e.g., object history), when compared to novel information (i.e., schematic and novel lures), and that this similarity will appear in the vmPFC, HC, and MTG regions. Yet we predict the opposite relationship in visual cortices, such that novelty irrespective of schematic association (e.g., both schematic and non-schematic lures) will show more similarity of neural patterns within the visual cortex, and in particular the early visual cortex which has been shown to be responsible for novelty and visual processing (Bowman & Dennis, 2015a; Kafkas & Montaldi, 2014; Koutstaal et al., 2001; Kurkela & Dennis, 2016; Slotnick & Schacter, 2006). This pattern of results would suggest that there is greater neural confusability (Simmonite & Polk, 2022) between schematic information in regions that process schema information, but that novel signals are more prominent in regions that process the perceptual properties of the novel items. Finally, relating neural similarity to behavior, we hypothesize that this similarity between schematic information (irrespective of object history) is what drives high hit and false alarm rates to schematic lures. Considering that previous work finds similar rates between hits and false alarms of schematic information (Lampinen et al., 2001; Miller & Gazzaniga, 1998; Webb et al., 2016), we propose that this neural similarity may account for these behavioral similarities.

## Materials & Methods

### Participants

Fifty-five right-handed native English speakers from the Penn State University and State College community completed this experiment as a part of a primary analysis of comparing younger and older adults, however due to the lack of age differences in the behavioral results, the current set of analyses collapses across both (see results for these statistics). Four participants were excluded from the analysis due to head motion in excess of 4 mm. Two were excluded for excessive atrophy. Seven additional participants were also excluded for poor behavioral performance (greater than 50% miss rate for schematic targets; three for a no response rate greater than 30%), leaving data from 42 participants reported in all analyses [29 females; mean age= 47.5 years (SD=26.65)]. A post hoc power analysis was run at the request of a reviewer, which suggested a power analysis of 42 participants was sufficient to reach a medium effect size (Cohen’s D= 0.5) and power of 89% using a one-sample t-test.

All participants provided written informed consent and received financial compensation for their participation. All experimental procedures were approved by The Pennsylvania State University’s Institutional Review Board for the ethical treatment of human participants.

### Stimuli and Task Procedure

The following information can also be found in Webb & Dennis (2016). Stimuli consisted of 26 schematic scenes (e.g., Bathroom, Farm), comprised of objects commonly associated with each schema (schematic targets: e.g., Farm: pig; Bathroom: toilet) as well as items unrelated to the schema (non-schematic targets: e.g., Farm: bush; Bathroom: vase). Lures consisted of both items commonly associated with each schema (schematic lures: e.g., Farm: tractor; Bathroom: sink), as well as non-schematic lures (e.g., piano, car, etc.) (See Fig 1.). All backgrounds and images were obtained from an internet image search. All items included in testing were normed for their association with each scene (See Webb and colleagues, 2016 for additional information). Each scene had an associated 4 schematic targets and 2-3 non-schematic targets at encoding. At retrieval, these 4 schematic targets, 2-3 non-schematic targets, 4 schematic lures were presented per scene in addition to 30 total non-schematic lures. (See Appendix A for full list of stimuli).

**Figure 1.**
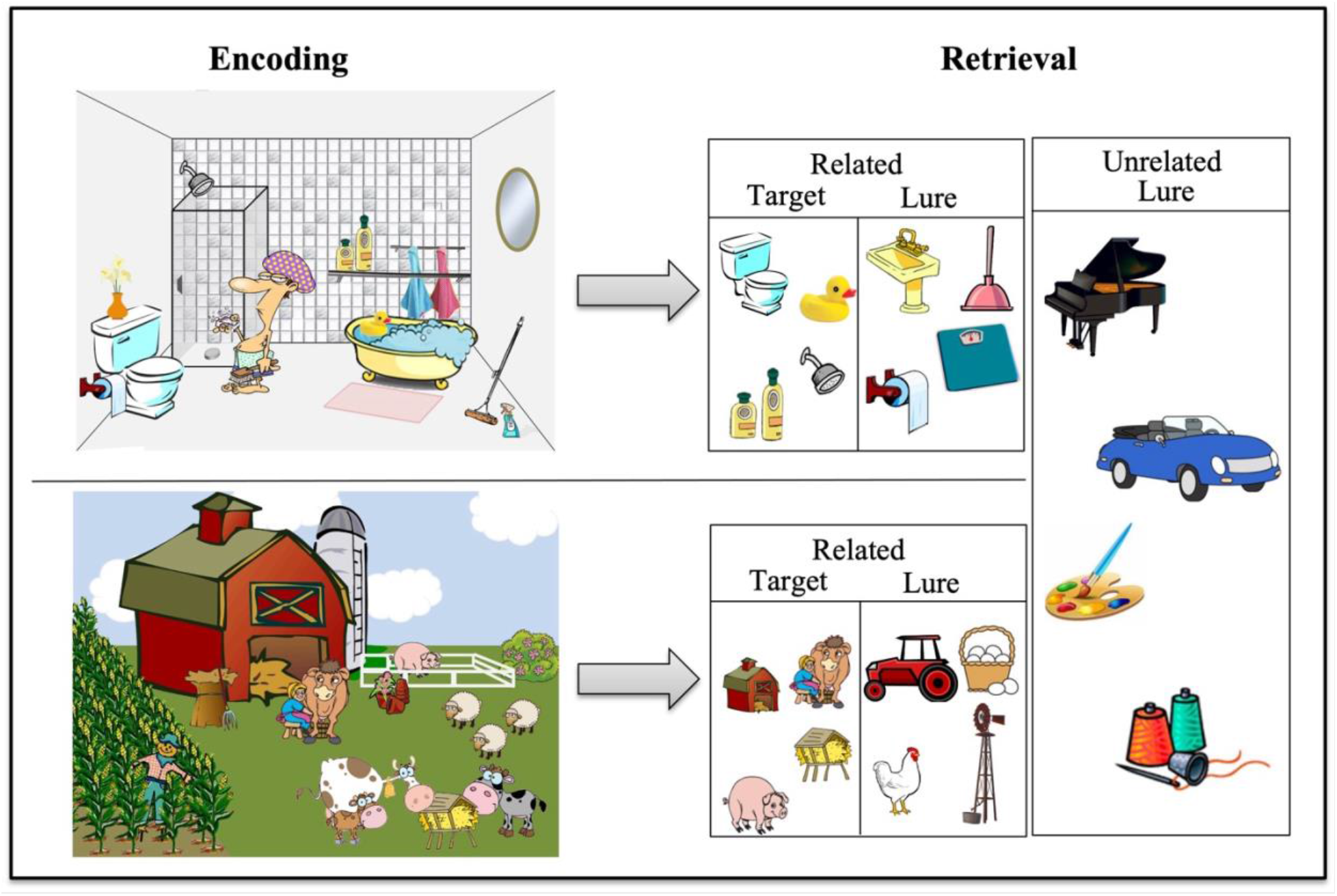
Task paradigm of two scenes seen at encoding and items seen at retrieval.

Encoding took place outside of the scanner while retrieval occurred in the scanner with approximately 30 minutes separating the two memory phases. Participants were asked to look at each scene and try to remember as much as they could for a later memory task. The 26 encoding scenes were presented for 10 seconds each across 2 runs, with 13 scenes presented in each run. During retrieval, all images (schematic targets from the scenes, schematic lures and non-schematic targets and lures) were presented in the center of the screen with three response options Remember, Know, New (RKN) displayed below each image. Participants completed 6 runs of approximately 7 minutes each in length at retrieval. Each image was displayed for 3 seconds. The images were pseudo-randomly sorted, ensuring that no more than 3 images from any one trial type appeared in a row and no 2 images associated with a given scene appeared in a row. In accordance with typical RKN task instructions, participants were told to respond ‘Remember’ if they could recollect specific details about the object such as its shape, color, placement in the scene or their thoughts or feelings during its initial presentation. Participants were told to respond ‘Know’ if the picture looked familiar, but they could not recollect any specific details of its prior presentation. They were told to respond ‘New’ if they believed the picture was not presented during the encoding session (see Figure 1). In total, there were 104 schematic targets, 104 schematic lures, 62 non-schematic targets and 30 non-schematic lures.

Images were projected onto a screen that participants viewed through a mirror attached to the head coil. Behavioral responses were recorded with the participant’s right hand on a 4-button response box. Images were displayed by COGENT in MATLAB (Mathworks Inc., Natick, MA, USA). Noise in the scanner was reduced with headphones, and cushioning was provided in the head coil to minimize head motion.

### Scanning parameters

Structural and functional images were acquired using a Siemens 3T scanner equipped with a 12-channel head coil, parallel to the AC–PC plane. Structural images were acquired with a 1,650 ms TR, 2.03 ms TE, 256 mm field of view (FOV), 256^2^ matrix, 160 axial slices, and 1.0 mm slice thickness for each participant. Echo-planar functional images were acquired using a descending acquisition, 2,500 ms TR, 25 ms TE, 240 mm FOV, an 80^2^ matrix, 90° flip angle, 42 axial slices with 3.0 mm slice thickness resulting in 3.0 mm isotropic voxels.

### MRI Data Preprocessing

*R*esults included in this manuscript come from preprocessing performed using fMRIPrep 20.1.1 (Esteban et al., 2019) which is based on Nipype 1.5.0 (Gorgolewski et al., 2011).

### Anatomical data preprocessing

A total of 1 T1-weighted (T1w) images were found within the input BIDS dataset. The T1-weighted (T1w) image was corrected for intensity non-uniformity (INU) with N4BiasFieldCorrection (Tustison et al., 2010), distributed with ANTs 2.2.0 (Avants et al., 2008), and used as T1w-reference throughout the workflow. The T1w-reference was then skull-stripped with a Nipype implementation of the antsBrainExtraction.sh workflow (from ANTs), using OASIS30ANTs as target template. Brain tissue segmentation of cerebrospinal fluid, white-matter and gray-matter was performed on the brain-extracted T1w using fast (FSL 5.0.9, Zhang et al., 2001). Brain surfaces were reconstructed using recon-all (FreeSurfer 6.0.1, Dale et al., 1999), and the brain mask estimated previously was refined with a custom variation of the method to reconcile ANTs-derived and FreeSurfer-derived segmentations of the cortical gray-matter of Mindboggle (Klein et al., 2017). Volume-based spatial normalization to one standard space (MNI152NLin2009cAsym) was performed through nonlinear registration with antsRegistration (ANTs 2.2.0), using brain-extracted versions of both T1w reference and the T1w template. The following template was selected for spatial normalization: ICBM 152 Nonlinear Asymmetrical template version 2009c (Fonov et al., 2009; TemplateFlow ID: MNI152NLin2009cAsym).

### Functional data preprocessing

For each of the 6 BOLD runs found per subject (across all tasks and sessions), the following preprocessing was performed. First, a reference volume and its skull-stripped version were generated using a custom methodology of fMRIPrep. Head-motion parameters with respect to the BOLD reference (transformation matrices, and six corresponding rotation and translation parameters) are estimated before any spatiotemporal filtering using mcflirt (FSL 5.0.9, Jenkinson et al., 2002). Susceptibility distortion correction (SDC) was omitted. The BOLD reference was then co-registered to the T1w reference using bbregister (FreeSurfer) which implements boundary-based registration (Greve & Fischl, 2009). Co-registration was configured with six degrees of freedom. The BOLD time-series (including slice-timing correction when applied) were resampled onto their original, native space by applying the transforms to correct for head-motion. These resampled BOLD time-series will be referred to as preprocessed BOLD in original space, or just preprocessed BOLD. The BOLD time-series were resampled into standard space, generating a preprocessed BOLD run in MNI152NLin2009cAsym space. First, a reference volume and its skull-stripped version were generated using a custom methodology of fMRIPrep. The head-motion estimates calculated in the correction step were also placed within the corresponding confounds file. The confound time series derived from head motion estimates and global signals were expanded with the inclusion of temporal derivatives and quadratic terms for each (Satterthwaite et al., 2013). Frames that exceeded a threshold of 0.5 mm FD or 1.5 standardized DVARS were annotated as motion outliers. All resamplings can be performed with a single interpolation step by composing all the pertinent transformations (i.e. head-motion transform matrices, susceptibility distortion correction when available, and co-registrations to anatomical and output spaces). Gridded (volumetric) resamplings were performed using antsApplyTransforms (ANTs), configured with Lanczos interpolation to minimize the smoothing effects of other kernels (Lanczos, 1964). Non-gridded (surface) resamplings were performed using mri_vol2surf (FreeSurfer). Many internal operations of fMRIPrep use Nilearn 0.6.2 (Abraham et al., 2014), mostly within the functional processing workflow. For more details of the pipeline, see <u>the section corresponding to</u> <u>workflows in *fMRIPrep*’s documentation</u>.

### Analyses

This study was pre-registered as a secondary analysis on a pre-existing data set on OSF: https://osf.io/mtd8r. This data has been published prior in a sample of younger adults (Webb et al., 2016) and in a sample of older adults (Webb & Dennis, 2019) examining univariate activation for schematic true and false memory.

### Behavioral Analyses

At the request of a reviewer and for transparency purposes, four mixed model ANOVAs were run (Age: young or old, condition: schematic or non-schematic images) predicting both recollected and adjusted familiarity hits and false alarm rates. Adjusted familiarity hits were calculated as pKnow Hits/(1 - pRemember Hits) and adjusted familiarity FA were calculated as pKnow FA/(1 - pRemember FA). These calculations take into account the fact the recollection and familiarity are not mutually exclusive processes (Duarte et al., 2006, 2010; Yonelinas, 2002; Yonelinas & Jacoby, 1995). Main effects and interactions were parsed out using paired t-tests. These were conducted using the rstatix package in R Studio (Kassambara, 2021).

### Pattern Similarity Analysis

To estimate neural activity associated with individual trials, separate GLMs were estimated in SPM12 defining one regressor for each trial at retrieval (Mumford et al., 2012). An additional six nuisance regressors were included in each run corresponding to motion. Whole-brain beta parameter maps were generated for each trial at retrieval for each participant. For any given parameter map, the value of each voxel represents the regression coefficient for that trial’s regressor in a multiple regression containing all other trials in the run and the motion parameters. These beta parameter maps were next concatenated across runs.

The main regions of interest include the hippocampus (HC), the middle occipital cortex (MOC), inferior occipital cortex (IOC), middle frontal gyrus (MFG), middle temporal gyrus (MTG) (Webb et al., 2016; Webb & Dennis, 2019) and the ventromedial prefrontal cortex (vmPFC ; van Kesteren et al., 2010; Warren et al., 2014). Activation in HC and vmPFC has been associated with schema-consistent information (Guo & Yang, 2020; van Kesteren et al., 2010; Warren et al., 2014). Additionally, visual cortex, including the IOC and MOC, is known for its ability to discriminate between old and new information (Bowman & Dennis, 2015b; Kurkela & Dennis, 2016; Slotnick & Schacter, 2006). Finally, activation in the MTG has been associated with false memories while the MFG has been implicated in both true and false memory (Webb et al., 2016). The ROIs were defined using anatomical masks of each bilateral region using the Wake Forest aal pickatlas in SPM12.

Two representational similarity analyses (RSA) were conducted to examine the overlap in representation of 1) schematic targets and schematic lures and 2) schematic lures and non-schematic lures at retrieval. Our analyses were aimed at identifying how object history (novel lure information) versus schematic membership (belonging to a schema) was represented neurally. The purpose of the RSA across schematic lures and non-schematic lure analysis was to identify regions that represented objective novelty across the two stimulus types (irrespective of schematic membership). Specifically, we aimed to explore to what extent neural patterns associated with schematic lures and non-schematic lures overlap. Additionally, the purpose of the RSA across schematic lures and schematic targets was to identify regions that represented schematic processing (irrespective of object history). Specifically, we aimed to explore to what extent neural patterns associated with schematic lures and schematic targets overlap.

Pattern analyses were conducted using the CoSMoMVPA toolbox (Oosterhof et al., 2016). Specifically, given our interest in understanding whether neural patterns seen with schematic lures are more similar to schematic target or non-schematic lure information, we compared the between-category similarity of lure information and schematic information irrespective of behavior. Additionally, we examined the between-category similarity of schematic hits and schematic false alarms to determine if increased pattern similarity is related to increases in false memories for schematic information. The similarities were directly tested via a paired t-test in RStudio. The similarity of schematic targets and lures will be referred to as similarity between schematic information. The similarity of schematic lures and non-schematic lures will be referred to as similarity between lure information. Significant results were confirmed via 10,000 permutation t-tests using the Mkinfer package in R (Kohl, 2022). Following this initial analysis interested in trials at the condition level, we did not have enough trial numbers of false alarms to non-related lures to behaviorally bin trials in the “lure similarity” analysis (see results for reporting of averages). This was a secondary data analysis, and the initial purpose of the study was not to examine non-schematic lures but rather focused on schematic targets and lures. Thus, the original design had fewer trial types in the design of the task.

### Exploratory RSA Searchlight

To investigate neural similarity in regions outside of our a priori ROIs, we executed the foregoing analysis using a whole brain mask of the cortex as an exploratory analysis. These analyses were conducted using the representational similarity analysis searchlight implemented in CoSMoMVPA toolbox (Oosterhof et al., 2016). Both analyses used a searchlight radius of 6 mm. The beta maps were then submitted to a paired t-test using SPM12 (http://fil.ion.ucl.ac.uk/spm/) to examine where similarity was greater for schematic information compared to lure information and vice versa. Due to the exploratory nature of this analysis, all second level results were thresholded at *p*<.001 using an extent threshold of 11 voxels based off AFNI’s 3dClustSim (Cox, 1996; Cox & Hyde, 1997).

## Results

### Behavioral Results

A 2x2 (age: old, young; Condition: schematic, non-schematic) mixed model ANOVA predicting recollected hits revealed no main effect of age (F(1,40) = 0.09, *p*=0.76, pes=.002), nor interaction between condition and age (F(1,40) =1.83, *p*=.18, pes=.044). A main effect of condition was revealed, F(1,40) = 253.36, *p*<.001, pes=.86, such that there were greater recollected hits to schematic targets (M=0.44, SD=0.13) compared to non-schematic targets (M=0.23, SD=0.12).

A 2x2 (age: old, young; Condition: schematic, non-schematic) mixed model ANOVA predicting adjusted familiarity hits revealed no main effect of age (F(1,40) =1.09. *p*=.30, pes=.026), nor interaction between condition and age (F(1,40) = 0.64, *p*=.43, pes=.016). A main effect of condition was revealed, F(1,40) =6.28, *p*=.016, pes=0.14, such that there were greater familiarity hits to schematic targets (M=0.46 SD=0.16) compared to non-schematic targets (M=0.42, SD=0.18).

A 2x2 (age: old, young; Condition: schematic, non-schematic) mixed model ANOVA predicting recollected FAs revealed that there was no main effect of age (F(1,40) = 0.10. *p*=.75, pes=.003), nor interaction between condition and age (F(1,40) = 0.43, *p*=.52, pes=.011). A main effect of condition was revealed, F(1,40) =94.82, *p*<.001, pes=.70, such that there were greater recollected FAs to schematic lures (M=0.23, SD=0.14) compared to non-schematic lures (M=0.07, SD=0.07).

A 2x2 (age: old, young; Condition: schematic, non-schematic) mixed model ANOVA predicting adjusted familiarity FAs revealed that there was no main effect of age (F(1,40) =0.12, *p*=.73, pes=.003), nor interaction between condition and age (F(1,40) = 0.005, *p*=.94, pes=.0001). A main effect of condition was revealed, F(1,40) =106.23, *p*<.001, pes=.72, such that there were greater adjusted familiarity FAs to schematic lures (M=0.41, SD=0.15) compared to non-schematic lures (M=0.20, SD=0.17).

As there were no main effects or interactions with age behaviorally, all neural analyses were collapsed across age.

### Pattern Similarity Results

Two RSAs were run and compared via a t-test. Schematic information refers to the RSA between schematic targets and schematic lures, while lure information refers to the RSA between schematic lures and non-schematic lures. There was significantly greater similarity between schematic information than lure information in the vmPFC, the MOC, and in the IOC, [*t*(41) = -2.40, *p*=0.021; *t*(41) = -5.21, *p*<.001; *t*(41) = -3.94, *p*<.001 respectively]. There were no significant differences between the neural similarity of schematic information and lure information in the MTG, MFG or HC (all *p*’s>.05) (see Table 1 for means). The current pattern of results indicates that overlapping neural patterns of schematic information, in the early and late visual cortices as well as the vmPFC, irrespective of whether subjects had seen it before or not, is more similar than information that is completely novel (see Figure 2 for RSA visualization).

**Figure 2.**
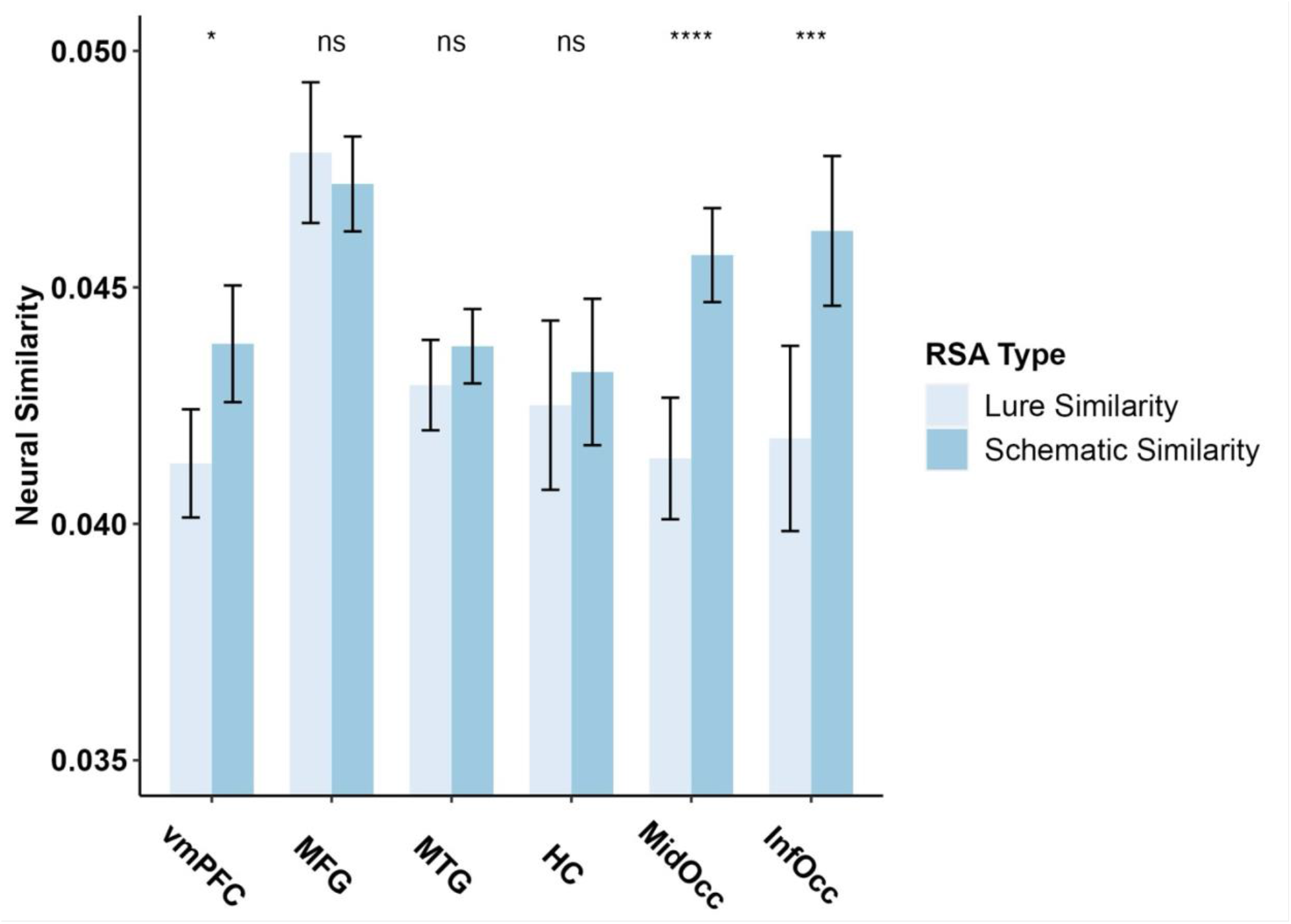
RSA Figure examining the difference between neural similarity of lures (novel lures and schematic lures) and schematic information (schematic targets and schematic lures). *Lure Similarity* = Similarity between Schematic Lures & Non-Schematic Lures; *Schematic Similarity*= Similarity between Schematic targets & Schematic Lures. ROI = region of interest.

**Table 1.** Means and standard deviation of pattern similarity between schematic information (schematic targets and lures) and lure information (schematic and novel lures).

|  | Schematic Similarity | Lure Similarity |
| --- | --- | --- |
|  | M(SD) | M(SD) |
| vmPFC* | 0.044 (.01) | 0.041 (.01) |
| MFG | 0.047 (.01) | 0.048 (.01) |
| MTG | 0.044 (.01) | 0.043 (.01) |
| HC | 0.043 (.01) | 0.043 (.01) |
| MOC**** | 0.046 (.01) | 0.041 (.01) |
| IOC*** | 0.046 (.01) | 0.042 (.01) |

### Exploratory RSA Searchlight

A contrast for similarity between lure information greater than similarity between schematic information (schematic lure & non-schematic lure similarity > schematic target & schematic lure similarity) of two searchlights conducted within a whole brain mask revealed no significant clusters. Additionally, a contrast for similarity between schematic information greater than similarity between lure information (schematic target & schematic lure similarity > schematic lure & non-schematic lure similarity) of two searchlights conducted within a whole brain mask revealed several significant clusters in the superior frontal gyrus and superior medial gyrus, HC, MFG and in the MTG (see Table 2 for full results and voxel locations and Figure 3 for visualization).

**Figure 3.**
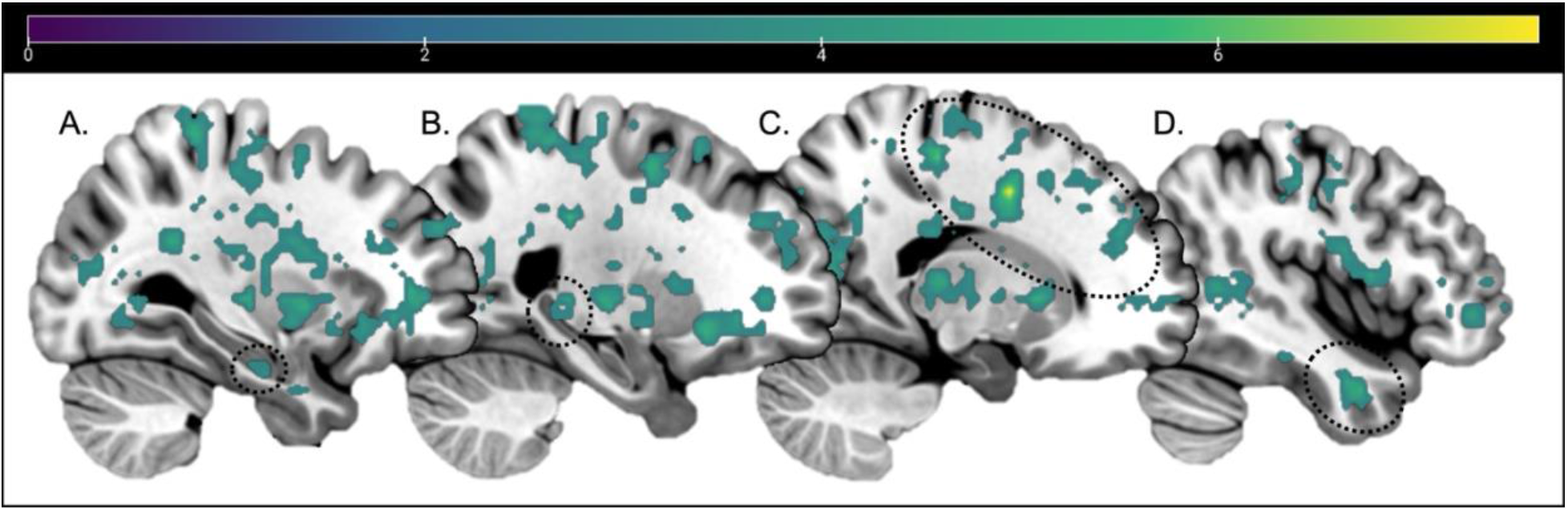
Whole Brain RSA searchlight contrast results for Schematic information similarity > Lure information similarity, thresholded at *p<*.001 and 11 voxels. No clusters survived thresholding for Lure information similarity > Schematic information similarity. Sagittal slices: - 30, -24, 17, 44. Dotted circles around: [A] Anterior HC, [B] Posterior HC, [C] MFG, [D] MTG. Color bar indicates t-values.

**Table 2.** RSA Searchlight results. PHG = parahippocampal gyrus; H= hemisphere; k = cluster threshold; t = statistical peak t-value; MNI x, y, and z coordinates; BA= Brodmann Area.

|  | BA | Hemisphere | k | t | x | y | z<br>{mm} |
| --- | --- | --- | --- | --- | --- | --- | --- |
| ***Schema > Lure Similarity*** |  |  |  |  |  |  |  |
| Superior-medial Frontal Gyrus | 10 | R | 16 | 4.51 | 12 | 60 | 18 |
| Ventral-medial Prefrontal Cortex | 10 | M | 47 | 4.87 | 8 | 56 | -4 |
|  | 11 | M | 11 | 4.65 | 8 | 24 | -18 |
| Middle Frontal Gyrus | 24 | R | 1477 | 8.58 | 18 | 2 | 42 |
| Precentral Gyrus | 44 | R |  | 6.85 | 50 | 6 | 8 |
|  | 9 | R | 14 | 4.65 | 36 | 38 | 36 |
|  | 6 | R | 16 | 4.05 | 18 | 6 | 56 |
| Superior Frontal Gyrus | - | L | 11 | 4.71 | -30 | 32 | 20 |
| Caudate | - | L | 14 | 5.17 | -12 | 12 | 0 |
| Middle Temporal Gyrus | 21 | R | 63 | 5.87 | 44 | 2 | -30 |
|  | 21 | L | 87 | 5.85 | -60 | -24 | -6 |
| Anterior PHG/Hippocampus | 20 | L | 37 | 5.2 | -40 | -10 | -30 |
| Posterior Hippocampus | - | L | 22 | 4.9 | -24 | -30 | 2 |
| Cingulum | 23 | M | 46 | 4.73 | -4 | -22 | 30 |
| Thalamus | - | R | 128 | 5.89 | 20 | -24 | 8 |
| Fusiform Gyrus | 20 | R | 31 | 4.66 | 42 | -28 | -16 |
| Paracentral Lobule | 4 | L | 1856 | 6.31 | -12 | -34 | 72 |
| Postcentral Gyrus | 6 | L |  | 6.29 | -54 | -4 | 42 |
| Putamen/Globus Pallidus | - | L |  | 6.28 | -22 | -12 | 2 |
| Cingulate | - | M | 11 | 4.59 | -4 | -46 | 6 |
| Precuneus | 31 | M | 13 | 5.23 | -6 | -64 | 36 |
| Occipital Gyrus | 19 | R | 147 | 5.79 | 36 | -54 | 2 |
|  | - | L | 171 | 5.75 | -34 | -66 | 8 |
| Middle Occipital Cortex | 18 | R | 265 | 6.4 | 18 | -84 | 0 |
| Inferior Occipital Cortex | 18 | L | 15 | 4.69 | -24 | -88 | -4 |
| ***Lure > Schema Similarity*** |  |  |  |  |  |  |  |
| *no significant clusters* |  |  |  |  |  |  |  |

### Pattern Similarity and Behavior

Since the above pattern similarity analyses were conducted on only target and lure information, we also wanted to directly determine whether neural pattern similarity between schematic hits and schematic false alarms was associated with increased hits and false alarms as well. Due to low false alarm numbers (avg = 1.98 FAs) in the non-schematic lure condition, we were unable to correlate or perform a powerful enough representational similarity analysis with behavior associated with this condition. Thus, we conducted an RSA to examine the neural similarity between recollected schematic hits and schematic false alarms to directly correlate to behavior. There were no significant correlations between pattern similarity for schematic recollected hits and schematic recollected false alarms with recollected hits or recollected false alarms (all *p*’s>.05).

## Discussion

The current study aimed to determine the neural mechanisms underlying false memories to schematic lures specifically in relation to schematic targets and non-schematic lures. In line with our predictions, several brain regions implicated in schematic processing, including the vmPFC and bilateral MTG, exhibited greater neural similarity regarding schematic content, irrespective of object history (i.e., schematic targets and schematic lures), as opposed to object novelty (i.e., schematic lures and unrelated lures). Contrary to our predictions, the same pattern of results was also observed throughout visual cortices, including both early and late visual cortices. In fact, no brain region demonstrated greater similarity in neural processing with respect to object novelty. This suggests that a relationship with previous schematic knowledge is more influential when processing novel information than object history itself. The results highlight the critical relationship between information belonging to a common schema and underscore how related information is processed in a manner that leads to its misidentification in memory tasks.

Greater neural pattern similarity for schematic targets and schematically related lures compared to that of both types of lures within the vmPFC and bilateral MTG is consistent with a larger literature implicating these regions in schematic processing. Specifically, the finding that the vmPFC exhibits greater neural similarity between schematic information expands upon prior work examining schemas at a univariate level (Guo & Yang, 2020; van Kesteren et al., 2010, 2013; Webb et al., 2016; Webb & Dennis, 2019) by informing us about the nature of the relationships between neural patterns in this region. Specifically, prior work has shown that the vmPFC is active during both the encoding and retrieval of schematic information (van Kesteren et al., 2010, 2013). The current work extends these findings suggesting that the schematic processing at retrieval underscoring targets is similar to that of schematic lures in this region, highlighting the broad nature of schema processing within the vmPFC. Given that there is greater neural similarity, and thus greater neural confusability, within this region for schematic information compared to novel information (Simmonite & Polk, 2022), this may also point to a common neural mechanism underlying high rates of both false alarms and hits to schematic information. However, as we did not find any correlations to behavior, future work is needed to confirm this relationship between pattern similarity and behavior.

While the MTG, MFG, and MTL ROIs as a whole did not exhibit significant differences in pattern similarity across trial types, an exploratory RSA searchlight identified greater neural similarity patterns for schematic information compared to lure information in clusters of voxels in all three regions. Specifically, greater schematic processing was found in the left and right MTG. This indicates that while ROI as a whole is not sensitive to differences across stimulus categories, specific components within the MTG represent schematic information more similarly than novel information. In addition to a similar pattern found in the vmPFC, results suggest that subcomponents of these regions process schematic information in a similar manner irrespective of the item’s history. With respect to the MTG, this finding also extends prior univariate work showing that the MTG is active during false memories, as well as work showing that activity in this region increases with respect to false memory rate (Dennis et al., 2007, 2008; Garoff-Eaton et al., 2006; Gilboa & Marlatte, 2017; Kubota et al., 2006; Slotnick & Schacter, 2006; Turney & Dennis, 2017; Webb et al., 2016; Webb & Dennis, 2019). The current findings suggest that similarity in schematic processing, including schematic gist related to lure information, underlies these past findings. Taken together, the findings across vmPFC and MTG support our hypotheses regarding greater pattern similarity in regions responsible for processing schematic information irrespective of item history.

The searchlight analysis also identified a large cluster of voxels within the right MFG, left posterior HC, and left anterior parahippocampal gyrus (PHG) as showing greater neural pattern similarity across schematic targets and schematic lures compared to both types of lures. Similar to the vmPFC and MTG, the MFG has been implicated in semantic processing (Binder et al., 2009; Kircher et al., 2001; Mummery et al., 2000; Noppeney & Price, 2002; Wise & Price, 2006) and the retrieval of semantic gist (Buckner & Petersen, 1996; Gabrieli et al., 1998; Mummery et al., 2000; Noppeney et al., 2007; Simons et al., 2005; Wise & Price, 2006). While these regions did not show significant similarity between schematic targets and schematic lures in the ROI similarity analysis, their presence in the whole brain searchlight analysis indicates that subcomponents of these regions are indeed more sensitive to schematic properties of stimuli compared to object novelty. These findings highlight the importance of taking into account both laterality and more focal regions when examining fine-grained differences in neural similarity related to memory processing. Specifically, subregions of the HC and MFG may be more influential in representing patterns of similarity compared to the region at large. Greater similarity across schematic stimuli, opposed to novelty, continues to highlight the ubiquitous nature of semantics in memory processing, with semantic and schematic information biasing the way novel stimuli not previously studied during encoding is processed at the time of retrieval. Interestingly, this bias also extends to retrieval-related representations within the MTL. The HC and PHG, are typically seen in the literature as being responsible for the retrieval of previously encoded memory traces. In the case of target items, this would include representing the schematic relationship associated with objects. Yet, for lures, while the objects encompass the same schematic gist, there is no encoded trace to be retrieved. Common processing across targets and lures in the MTL may represent retrieval of this shared schematic gist incorrectly bound to the representation of the lure object. This error in neural processing within the MTL may be a critical factor in the lure item being incorrectly identified as “old”, leading to the high rate of false memories in schematic-based memory paradigms.

While overlapping schematic representations and gist processing was expected for targets and lures that are related to the encoding schema within regions previously implicated in schematic processing, it was expected that neural patterns in visual cortices would exhibit a dissociation between the two trial types. Specifically, given the absence of perceptual overlap between schematic lures and schematic targets, we expected visual processing regions to represent all novel information more similarly than the schematic information. Returning to the example in the introduction of the bathroom sink, this sink lure shares no perceptual features that are seen in target items from the encoded schema, such as a toilet or bathtub. This novelty of item identity and perceptual properties should be beneficial to memory processing and elicit unique processing within the visual cortices that would allow for the objects’ correct rejection in memory testing (Bowman & Dennis, 2015a, 2015b). However, both the IOC and MOC ROIs showed greater pattern similarity for schematic information irrespective of object history: the same pattern as observed in the vmPFC and MTG.

Prior univariate analyses conducted on this data found that visual regions were capable of distinguishing between true and false recollection of schematic information (Webb et al., 2016). However, the current pattern similarity analyses suggest that, despite this difference, neural patterns in visual cortices are more closely related for objects (irrespective of whether it was a target or lure) that share a common schema than for objects sharing a common presentation history. While contrary to our predictions, this is interesting regarding the often observed high false alarm rate in schematic memory studies. Results suggest that not only is a schema signal coming from regions such as vmPFC and MTG, which are known for processing relatedness amongst schematic items (van Kesteren et al., 2010, 2013; Webb et al., 2016; Webb & Dennis, 2019), but also from regions that are known for processing more discrete object properties such as the early and late visual cortices (Bowman & Dennis, 2015a; Gutchess & Schacter, 2012; Slotnick & Schacter, 2004, 2006). This level of similarity may also underlie greater confusability across targets and schematic lures, leading to both being identified as “old” at high rates during memory responses. That is, the neural patterns may be too similar across schematic targets and lures to correctly reject the related lure even though the lure is completely new and not part of the studied set. Previous work asserts that object history is critical to activation in visual regions, as well as critical to novelty detection (Bowman & Dennis, 2015a; Gutchess & Schacter, 2012; Slotnick & Schacter, 2004, 2006). However, the current results expand the nuance surrounding this idea, suggesting that information belonging to a prior schema may be more influential than object history with respect to processing in visual cortices (and throughout the brain), leading to its persuasive influence on memory processing over and above object novelty.

Finally, compared to neural pattern similarity for schematic information, lure similarity showed no significant clusters in the similarity searchlight. That is, no region, either within our *a-priori* ROIs nor through our searchlight analysis, exhibited greater neural similarity for item novelty greater than schematic similarity. This indicates that there is widespread similarity across the brain for schematic information, which overrides object history in terms of pattern similarity. This again highlights the pervasive influence of schemas throughout the brain in regions apart from those hypothesized initially that is not the case for novel information. While not directly related to behavior, these results suggest a mechanism behind why false memories to schematic information are so prevalent compared to that of novel information. This also replicates our ROI-based pattern similarity analyses that suggest that there is enhanced similarity amongst schematic information and thus dissimilarity amongst lure information in comparison.

Interestingly, despite the neural similarity for schematic information within cortical and subcortical regions, neural processing leads to the identification of novel information even when there is overlap in schematic relationships and neural signals between targets and lures. Thus, despite this similarity, this confusability can be overridden to an extent to allow for successful memory performance. However, absent of any region showing greater pattern similarity for item novelty than schematic similarity, this suggests that while schematic similarity *can* be overcome during memory retrieval (e.g., correct rejections to schematic lures), this may be a particularly difficult process given the neural bias related to the processing of schematic content when viewing objects part of a known schematic category of stimuli. Combined with the fact that participants also exhibit higher hit rates to schematic targets compared to non-schematic targets (Lampinen et al., 2001; Miller & Gazzaniga, 1998; Webb et al., 2016; Webb & Dennis, 2019), these findings support the prevalent nature of schemas in memory processing. However, absent of correlations with behavior, further work is needed to directly link schematic processing with memory performance across both successful and unsuccessful memory responses.

Taken together, the current results suggest that previous knowledge of a certain schematic event has a greater impact on neural processing than the novelty of information across multiple brain regions including prefrontal, MTL, and visual cortices. Specifically, these and many other brain regions represent schematic information across items more similarly than object history. Most interestingly, this similarity of processing was found not only within regions known for schema processing, but throughout the brain, including clusters of voxels within the MTL and occipital cortices which have been traditionally associated with memory success. This similarity in neural processing across targets and lures may be why people false alarm at higher rates to schematic lures compared to non-schematic lures; however, given our absence of brain-behavior correlations and low false alarm numbers regarding the non-schematic trials, this latter point would need further work to verify. Finally, no brain region exhibited higher neural pattern similarity as a function of item history, suggesting that schematic processing is a strong predictor of neural activity above and beyond object novelty.

## Acknowledgements

This work was supported by a National Science Foundation Grant (BCS1025709) awarded to NAD. CMC was supported by National Institute on Aging Grant T32 AG049676 to The Pennsylvania State University. We thank Christina Webb and Indira Turney for past work collecting and processing this dataset. We also thank Alexa Becker, Becky Wagner and Luke Dubec for comments on previous drafts of this manuscript.

## Data Availability Statement

Data may be made available upon request of the corresponding author.

## Appendix A.

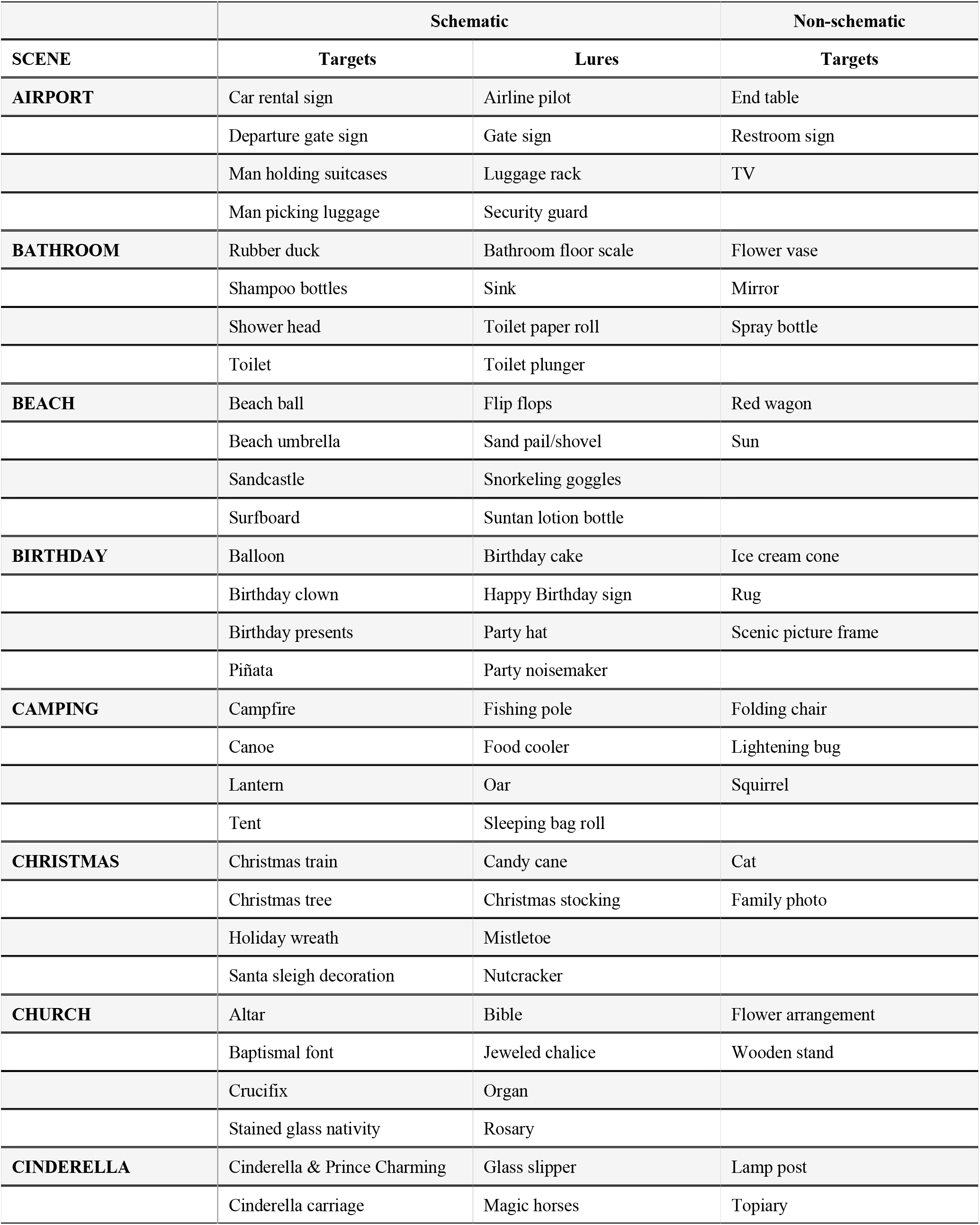
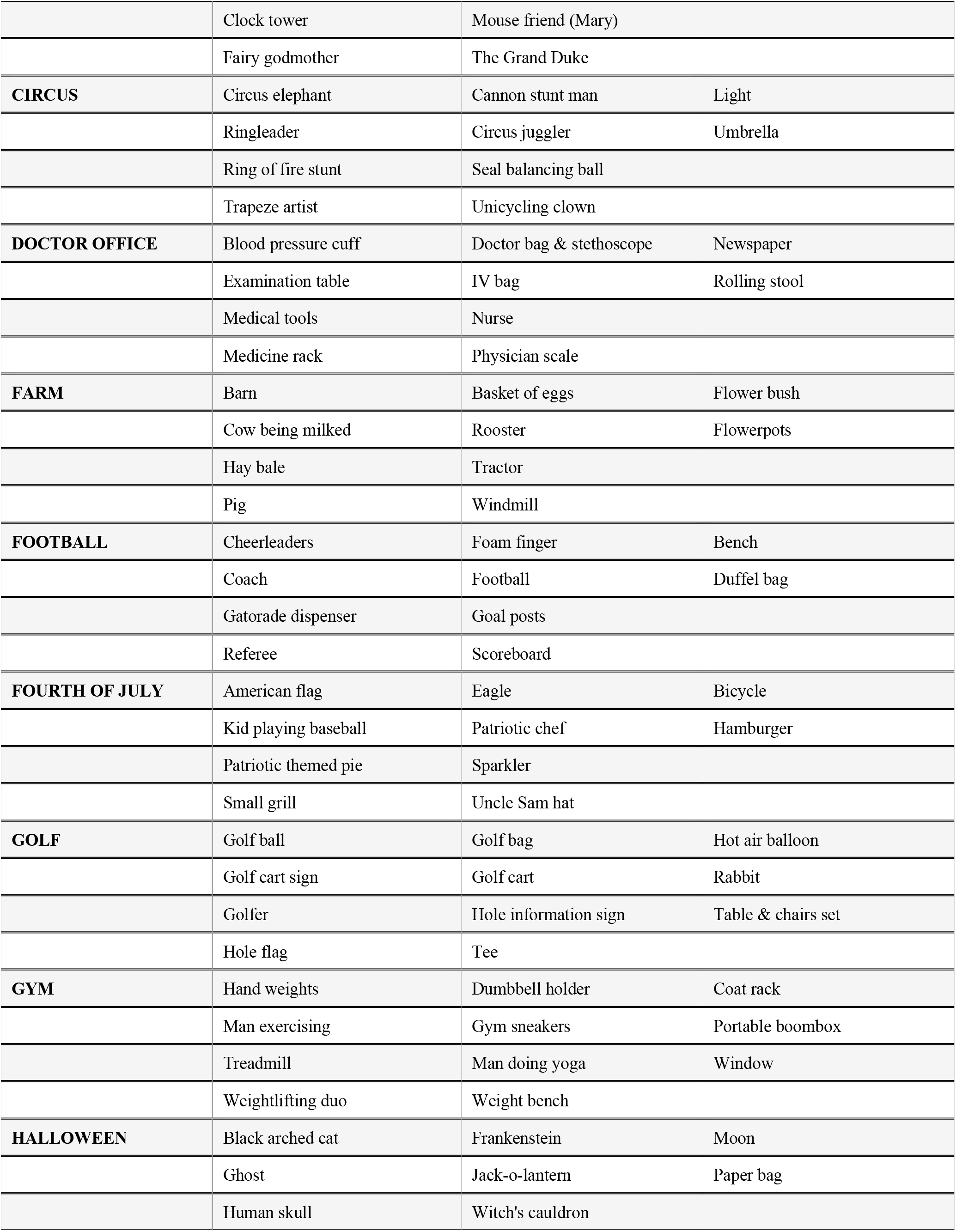
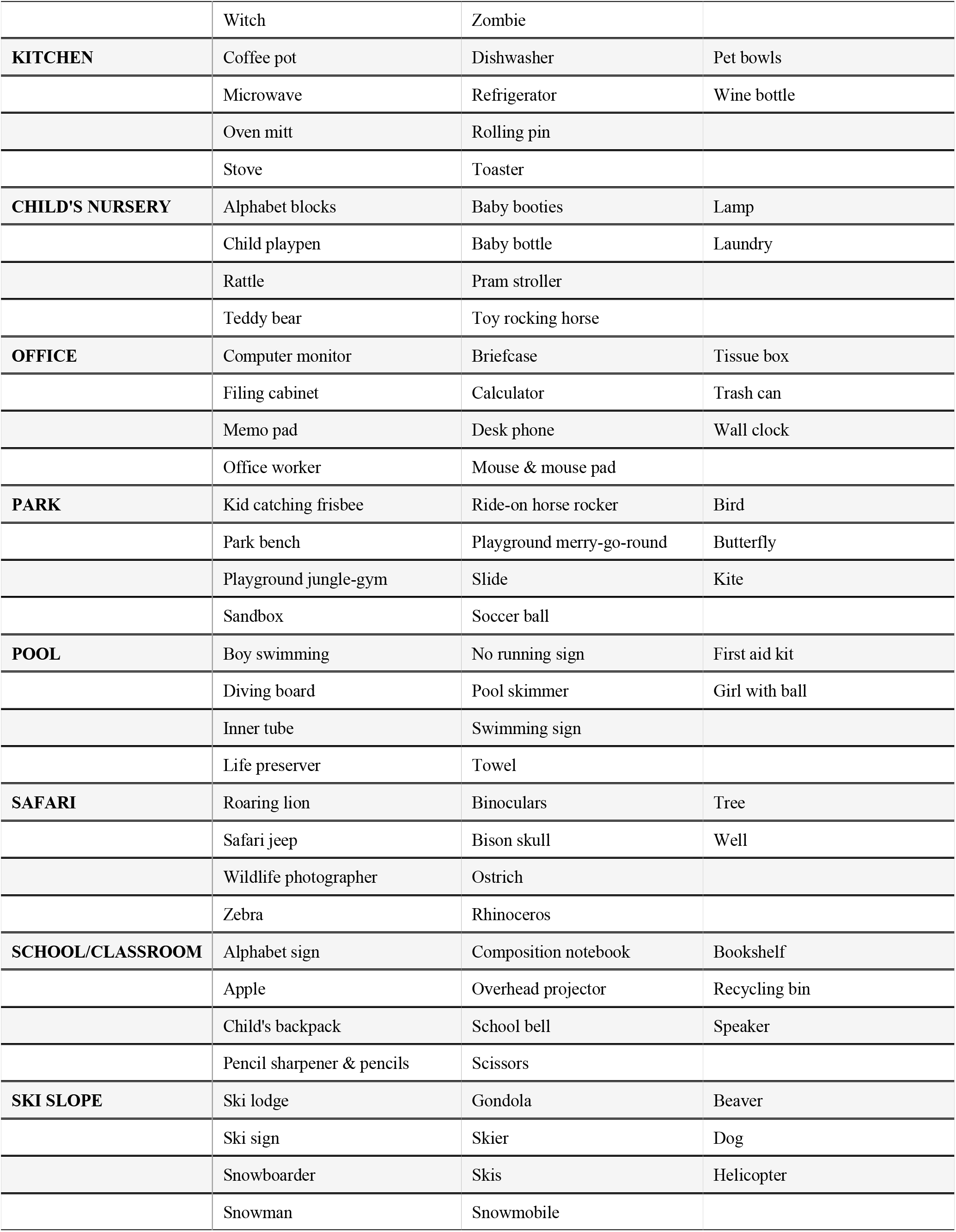
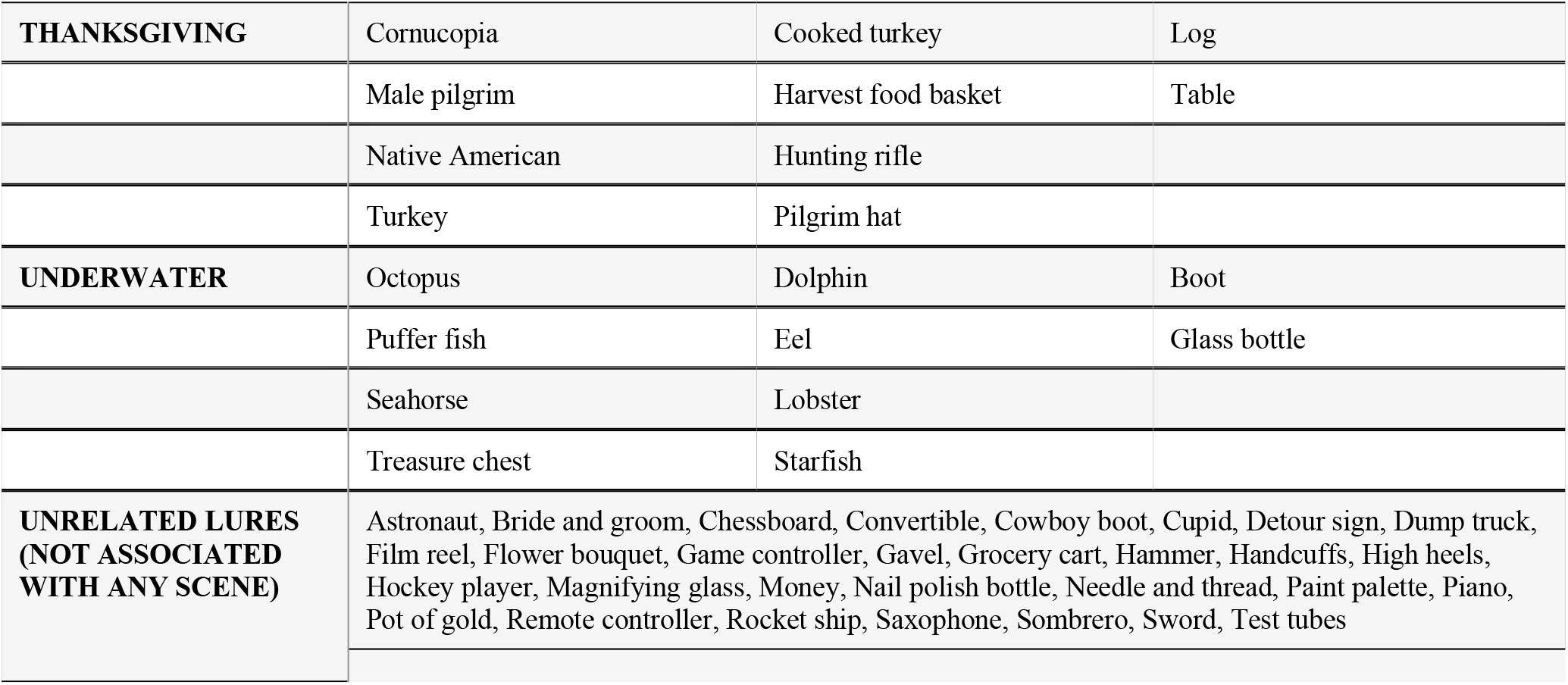

|  | Schematic |  | Non-schematic |
| --- | --- | --- | --- |
| SCENE | Targets | Lures | Targets |
| AIRPORT | Car rental sign | Airline pilot | End table |
|  | Departure gate sign | Gate sign | Restroom sign |
|  | Man holding suitcases | Luggage rack | TV |
|  | Man picking luggage | Security guard |  |
| BATHROOM | Rubber duck | Bathroom floor scale | Flower vase |
|  | Shampoo bottles | Sink | Mirror |
|  | Shower head | Toilet paper roll | Spray bottle |
|  | Toilet | Toilet plunger |  |
| BEACH | Beach ball | Flip flops | Red wagon |
|  | Beach umbrella | Sand pail/shovel | Sun |
|  | Sandcastle | Snorkeling goggles |  |
|  | Surfboard | Suntan lotion bottle |  |
| BIRTHDAY | Balloon | Birthday cake | Ice cream cone |
|  | Birthday clown | Happy Birthday sign | Rug |
|  | Birthday presents | Party hat | Scenic picture frame |
|  | Piñata | Party noisemaker |  |
| CAMPING | Campfire | Fishing pole | Folding chair |
|  | Canoe | Food cooler | Lightening bug |
|  | Lantern | Oar | Squirrel |
|  | Tent | Sleeping bag roll |  |
| CHRISTMAS | Christmas train | Candy cane | Cat |
|  | Christmas tree | Christmas stocking | Family photo |
|  | Holiday wreath | Mistletoe |  |
|  | Santa sleigh decoration | Nutcracker |  |
| CHURCH | Altar | Bible | Flower arrangement |
|  | Baptismal font | Jeweled chalice | Wooden stand |
|  | Crucifix | Organ |  |
|  | Stained glass nativity | Rosary |  |
| CINDERELLA | Cinderella & Prince Charming | Glass slipper | Lamp post |
|  | Cinderella carriage | Magic horses | Topiary |
|  | Clock tower | Mouse friend (Mary) |  |
|  | Fairy godmother | The Grand Duke |  |
| **CIRCUS** | Circus elephant | Cannon stunt man | Light |
|  | Ringleader | Circus juggler | Umbrella |
|  | Ring of fire stunt | Seal balancing ball |  |
|  | Trapeze artist | Unicycling clown |  |
| **DOCTOR OFFICE** | Blood pressure cuff | Doctor bag & stethoscope | Newspaper |
|  | Examination table | IV bag | Rolling stool |
|  | Medical tools | Nurse |  |
|  | Medicine rack | Physician scale |  |
| **FARM** | Barn | Basket of eggs | Flower bush |
|  | Cow being milked | Rooster | Flowerpots |
|  | Hay bale | Tractor |  |
|  | Pig | Windmill |  |
| **FOOTBALL** | Cheerleaders | Foam finger | Bench |
|  | Coach | Football | Duffel bag |
|  | Gatorade dispenser | Goal posts |  |
|  | Referee | Scoreboard |  |
| **FOURTH OF JULY** | American flag | Eagle | Bicycle |
|  | Kid playing baseball | Patriotic chef | Hamburger |
|  | Patriotic themed pie | Sparkler |  |
|  | Small grill | Uncle Sam hat |  |
| **GOLF** | Golf ball | Golf bag | Hot air balloon |
|  | Golf cart sign | Golf cart | Rabbit |
|  | Golfer | Hole information sign | Table & chairs set |
|  | Hole flag | Tee |  |
| **GYM** | Hand weights | Dumbbell holder | Coat rack |
|  | Man exercising | Gym sneakers | Portable boombox |
|  | Treadmill | Man doing yoga | Window |
|  | Weightlifting duo | Weight bench |  |
| **HALLOWEEN** | Black arched cat | Frankenstein | Moon |
|  | Ghost | Jack-o-lantern | Paper bag |
|  | Human skull | Witch's cauldron |  |
|  | Witch | Zombie |  |
| **KITCHEN** | Coffee pot | Dishwasher | Pet bowls |
|  | Microwave | Refrigerator | Wine bottle |
|  | Oven mitt | Rolling pin |  |
|  | Stove | Toaster |  |
| **CHILD'S NURSERY** | Alphabet blocks | Baby booties | Lamp |
|  | Child playpen | Baby bottle | Laundry |
|  | Rattle | Pram stroller |  |
|  | Teddy bear | Toy rocking horse |  |
| **OFFICE** | Computer monitor | Briefcase | Tissue box |
|  | Filing cabinet | Calculator | Trash can |
|  | Memo pad | Desk phone | Wall clock |
|  | Office worker | Mouse & mouse pad |  |
| **PARK** | Kid catching frisbee | Ride-on horse rocker | Bird |
|  | Park bench | Playground merry-go-round | Butterfly |
|  | Playground jungle-gym | Slide | Kite |
|  | Sandbox | Soccer ball |  |
| **POOL** | Boy swimming | No running sign | First aid kit |
|  | Diving board | Pool skimmer | Girl with ball |
|  | Inner tube | Swimming sign |  |
|  | Life preserver | Towel |  |
| **SAFARI** | Roaring lion | Binoculars | Tree |
|  | Safari jeep | Bison skull | Well |
|  | Wildlife photographer | Ostrich |  |
|  | Zebra | Rhinoceros |  |
| **SCHOOL/CLASSROOM** | Alphabet sign | Composition notebook | Bookshelf |
|  | Apple | Overhead projector | Recycling bin |
|  | Child's backpack | School bell | Speaker |
|  | Pencil sharpener & pencils | Scissors |  |
| **SKI SLOPE** | Ski lodge | Gondola | Beaver |
|  | Ski sign | Skier | Dog |
|  | Snowboarder | Skis | Helicopter |
|  | Snowman | Snowmobile |  |
| THANKSGIVING | Cornucopia | Cooked turkey | Log |
|  | Male pilgrim | Harvest food basket | Table |
|  | Native American | Hunting rifle |  |
|  | Turkey | Pilgrim hat |  |
| UNDERWATER | Octopus | Dolphin | Boot |
|  | Puffer fish | Eel | Glass bottle |
|  | Seahorse | Lobster |  |
|  | Treasure chest | Starfish |  |
| UNRELATED LURES<br>(NOT ASSOCIATED<br>WITH ANY SCENE) | Astronaut, Bride and groom, Chessboard, Convertible, Cowboy boot, Cupid, Detour sign, Dump truck, Film reel, Flower bouquet, Game controller, Gavel, Grocery cart, Hammer, Handcuffs, High heels, Hockey player, Magnifying glass, Money, Nail polish bottle, Needle and thread, Paint palette, Piano, Pot of gold, Remote controller, Rocket ship, Saxophone, Sombrero, Sword, Test tubes |  |  |

## References

Abe, N., Okuda, J., Suzuki, M., Sasaki, H., Matsuda, T., Mori, E., Tsukada, M., & Fujii, T. (2008). Neural Correlates of True Memory, False Memory, and Deception. Cerebral Cortex, 18(12), 2811–2819. 10.1093/cercor/bhn037

Abraham, A., Pedregosa, F., Eickenberg, M., Gervais, P., Mueller, A., Kossaifi, J., Gramfort, A., Thirion, B., & Varoquaux, G. (2014). Machine learning for neuroimaging with scikit-learn. Frontiers in Neuroinformatics, 8. https://www.frontiersin.org/articles/10.3389/fninf.2014.00014

Alba, J. W., & Hasher, L. (1983). Is memory schematic? Psychological Bulletin, 93, 203–231. 10.1037/0033-2909.93.2.203

Alboukadel Kassambara. (2021). *Rstatix: Pipe-Friendly Framework for Basic Statistical Tests. R package version 0.7.0*. https://CRAN.R-project.org/package=rstatix [Computer software].

Avants, B. B., Epstein, C. L., Grossman, M., & Gee, J. C. (2008). Symmetric diffeomorphic image registration with cross-correlation: Evaluating automated labeling of elderly and neurodegenerative brain. Medical Image Analysis, 12(1), 26–41. 10.1016/j.media.2007.06.004

Binder, J. R., Desai, R. H., Graves, W. W., & Conant, L. L. (2009). Where Is the Semantic System? A Critical Review and Meta-Analysis of 120 Functional Neuroimaging Studies. Cerebral Cortex (New York, NY), 19(12), 2767–2796. 10.1093/cercor/bhp055

Bowman, C. R., & Dennis, N. A. (2015a). Age differences in the neural correlates of novelty processing: The effects of item-relatedness. Brain Research, 1612, 2–15. 10.1016/j.brainres.2014.08.006

Bowman, C. R., & Dennis, N. A. (2015b). The neural correlates of correctly rejecting lures during memory retrieval: The role of item relatedness. Experimental Brain Research, 233(6), 1963–1975. 10.1007/s00221-015-4268-y

Brewer, W. F., & Treyens, J. C. (1981). Role of schemata in memory for places. Cognitive Psychology, 13(2), 207–230. 10.1016/0010-0285(81)90008-6

Brod, G., Lindenberger, U., & Shing, Y. L. (2017). Neural activation patterns during retrieval of schema-related memories: Differences and commonalities between children and adults. Developmental Science, 20(6), e12475. 10.1111/desc.12475

Buckner, R. L., & Petersen, S. E. (1996). What does neuroimaging tell us about the role of prefrontal cortex in memory retrieval? Seminars in Neuroscience, 8(1), 47–55. 10.1006/smns.1996.0007

Castel, A. D. (2005). Memory for grocery prices in younger and older adults: The role of schematic support. Psychology and Aging, 20, 718–721. 10.1037/0882-7974.20.4.718

Charlton, S. G., & Leov, J. (2021). Driving without memory: The strength of schema-consistent false memories. Transportation Research Part F: Traffic Psychology and Behaviour, 83, 12–21. 10.1016/j.trf.2021.09.018

Coane, J. H., McBride, D. M., Huff, M. J., Chang, K., Marsh, E. M., & Smith, K. A. (2021). Manipulations of List Type in the DRM Paradigm: A Review of How Structural and Conceptual Similarity Affect False Memory. Frontiers in Psychology, 12, 668550. 10.3389/fpsyg.2021.668550

Cox, R. W. (1996). AFNI: Software for analysis and visualization of functional magnetic resonance neuroimages. Computers and Biomedical Research, an International Journal, 29(3), 162–173. 10.1006/cbmr.1996.0014

Cox, R. W., & Hyde, J. S. (1997). Software tools for analysis and visualization of fMRI data. NMR in Biomedicine, 10(4–5), 171–178. 10.1002/(SICI)1099-1492(199706/08)10:4/5<171::AID-NBM453>3.0.CO;2-L

Dale, A. M., Fischl, B., & Sereno, M. I. (1999). Cortical Surface-Based Analysis: I. Segmentation and Surface Reconstruction. NeuroImage, 9(2), 179–194. 10.1006/nimg.1998.0395

Dennis, N. A., Bowman, C. K., & Turney, I. C. (2015). Functional neuroimaging of false memories. In The Wiley handbook on the cognitive neuroscience of memory (pp. 150– 171). Wiley Blackwell. 10.1002/9781118332634.ch8

Dennis, N. A., Kim, H., & Cabeza, R. (2007). Effects of aging on true and false memory formation: An fMRI study. Neuropsychologia, 45(14), 3157–3166. 10.1016/j.neuropsychologia.2007.07.003

Dennis, N. A., Kim, H., & Cabeza, R. (2008). Age-related Differences in Brain Activity during True and False Memory Retrieval. Journal of Cognitive Neuroscience, 20(8), 1390–1402. 10.1162/jocn.2008.20096

Dennis, N. A., & Turney, I. C. (2018). The influence of perceptual similarity and individual differences on false memories in aging. Neurobiology of Aging, 62, 221–230. 10.1016/j.neurobiolaging.2017.10.020

Duarte, A., Graham, K. S., & Henson, R. N. (2010). Age-related changes in neural activity associated with familiarity, recollection and false recognition. Neurobiology of Aging, 31(10), 1814–1830. 10.1016/j.neurobiolaging.2008.09.014

Duarte, A., Ranganath, C., Trujillo, C., & Knight, R. T. (2006). Intact recollection memory in high-performing older adults: ERP and behavioral evidence. Journal of Cognitive Neuroscience, 18(1), 33–47. 10.1162/089892906775249988

Esteban, O., Markiewicz, C. J., Blair, R. W., Moodie, C. A., Isik, A. I., Erramuzpe, A., Kent, J. D., Goncalves, M., DuPre, E., Snyder, M., Oya, H., Ghosh, S. S., Wright, J., Durnez, J., Poldrack, R. A., & Gorgolewski, K. J. (2019). fMRIPrep: A robust preprocessing pipeline for functional MRI. Nature Methods, 16(1), Article 1. 10.1038/s41592-018-0235-4

Fonov, V., Evans, A., McKinstry, R., Almli, C., & Collins, D. (2009). Unbiased nonlinear average age-appropriate brain templates from birth to adulthood. NeuroImage, 47, S102. 10.1016/S1053-8119(09)70884-5

Gabrieli, J. D. E., Poldrack, R. A., & Desmond, J. E. (1998). The role of left prefrontal cortex in language and memory. Proceedings of the National Academy of Sciences, 95(3), 906– 913. 10.1073/pnas.95.3.906

Garoff-Eaton, R. J., Kensinger, E. A., & Schacter, D. L. (2007). The neural correlates of conceptual and perceptual false recognition. Learning & Memory, 14(10), 684–692. 10.1101/lm.695707

Garoff-Eaton, R. J., Slotnick, S. D., & Schacter, D. L. (2006). Not All False Memories Are Created Equal: The Neural Basis of False Recognition. Cerebral Cortex, 16(11), 1645– 1652. 10.1093/cercor/bhj101

Gilboa, A., & Marlatte, H. (2017). Neurobiology of Schemas and Schema-Mediated Memory. Trends in Cognitive Sciences, 21(8), 618–631. 10.1016/j.tics.2017.04.013

Gorgolewski, K., Burns, C., Madison, C., Clark, D., Halchenko, Y., Waskom, M., & Ghosh, S. (2011). Nipype: A Flexible, Lightweight and Extensible Neuroimaging Data Processing Framework in Python. Frontiers in Neuroinformatics, 5. https://www.frontiersin.org/articles/10.3389/fninf.2011.00013

Greve, D. N., & Fischl, B. (2009). Accurate and robust brain image alignment using boundary-based registration. NeuroImage, 48(1), 63–72. 10.1016/j.neuroimage.2009.06.060

Guo, D., & Yang, J. (2020). Interplay of the long axis of the hippocampus and ventromedial prefrontal cortex in schema-related memory retrieval. Hippocampus, 30(3), 263–277. 10.1002/hipo.23154

Gutchess, A. H., & Schacter, D. L. (2012). The neural correlates of gist-based true and false recognition. NeuroImage, 59(4), 3418–3426. 10.1016/j.neuroimage.2011.11.078

Haxby, J. V., Gobbini, M. I., Furey, M. L., Ishai, A., Schouten, J. L., & Pietrini, P. (2001). Distributed and Overlapping Representations of Faces and Objects in Ventral Temporal Cortex. Science, 293(5539), 2425–2430. 10.1126/science.1063736

Jenkinson, M., Bannister, P., Brady, M., & Smith, S. (2002). Improved Optimization for the Robust and Accurate Linear Registration and Motion Correction of Brain Images. NeuroImage, 17(2), 825–841. 10.1006/nimg.2002.1132

Kafkas, A., & Montaldi, D. (2014). Two separate, but interacting, neural systems for familiarity and novelty detection: A dual-route mechanism. Hippocampus, 24(5), 516–527. 10.1002/hipo.22241

Kircher, T. T. J., Brammer, M., Tous Andreu, N., Williams, S. C. R., & McGuire, P. K. (2001). Engagement of right temporal cortex during processing of linguistic context. Neuropsychologia, 39(8), 798–809. 10.1016/S0028-3932(01)00014-8

Kleider, H. M., Pezdek, K., Goldinger, S. D., & Kirk, A. (2008). Schema-driven source misattribution errors: Remembering the expected from a witnessed event. Applied Cognitive Psychology, 22(1), 1–20. 10.1002/acp.1361

Klein, A., Ghosh, S. S., Bao, F. S., Giard, J., Häme, Y., Stavsky, E., Lee, N., Rossa, B., Reuter, M., Neto, E. C., & Keshavan, A. (2017). Mindboggling morphometry of human brains. PLOS Computational Biology, 13(2), e1005350. 10.1371/journal.pcbi.1005350

Kohl, M. (2022). *MKinfer: Inferential Statistics.* (R package version 0.7) [Computer software]. https://www.stamats.de.

Koutstaal, W., Wagner, A. D., Rotte, M., Maril, A., Buckner, R. L., & Schacter, D. L. (2001). Perceptual specificity in visual object priming: Functional magnetic resonance imaging evidence for a laterality difference in fusiform cortex. Neuropsychologia, 39(2), 184–199. 10.1016/s0028-3932(00)00087-7

Kubota, Y., Toichi, M., Shimizu, M., Mason, R. A., Findling, R. L., Yamamoto, K., & Calabrese, J. R. (2006). Prefrontal hemodynamic activity predicts false memory—A near-infrared spectroscopy study. NeuroImage, 31(4), 1783–1789. 10.1016/j.neuroimage.2006.02.003

Kurkela, K. A., & Dennis, N. A. (2016). Event-related fMRI studies of false memory: An Activation Likelihood Estimation meta-analysis. Neuropsychologia, 81, 149–167. 10.1016/j.neuropsychologia.2015.12.006

Lampinen, J., Copeland, S., & Neuschatz, J. (2001). Recollections of Things Schematic: Room Schemas Revisited. Journal of Experimental Psychology. Learning, Memory, and Cognition, 27, 1211–1222. 10.1037//0278-7393.27.5.1211

Lampinen, J. M., Faries, J. M., Neuschatz, J. S., & Toglia, M. P. (2000). Recollections of things schematic: The influence of scripts on recollective experience. Applied Cognitive Psychology, 14(6), 543–554. 10.1002/1099-0720(200011/12)14:6<543::AID-ACP674>3.0.CO;2-K

Lanczos, C. (1964). Evaluation of Noisy Data. *Journal of the Society for Industrial and Applied Mathematics: Series B*, Numerical Analysis, 1, 76–85.

Lew, A. R., & Howe, M. L. (2017). Out of place, out of mind: Schema-driven false memory effects for object-location bindings. *Journal of Experimental Psychology: Learning*, Memory, and Cognition, 43, 404–421. 10.1037/xlm0000317

Miller, M. B., & Gazzaniga, M. S. (1998). Creating false memories for visual scenes. Neuropsychologia, 36(6), 513–520. 10.1016/S0028-3932(97)00148-6

Mumford, J. A., Turner, B. O., Ashby, F. G., & Poldrack, R. A. (2012). Deconvolving BOLD activation in event-related designs for multivoxel pattern classification analyses. NeuroImage, 59(3), 2636–2643. 10.1016/j.neuroimage.2011.08.076

Mummery, C. J., Patterson, K., Price, C. J., Ashburner, J., Frackowiak, R. S. J., & Hodges, J. R. (2000). A voxel-based morphometry study of semantic dementia: Relationship between temporal lobe atrophy and semantic memory. Annals of Neurology, 47(1), 36–45. 10.1002/1531-8249(200001)47:1<36::AID-ANA8>3.0.CO;2-L

Neuschatz, J. S., Lampinen, J. M., Preston, E. L., Hawkins, E. R., & Toglia, M. P. (2002). The effect of memory schemata on memory and the phenomenological experience of naturalistic situations. Applied Cognitive Psychology, 16(6), 687–708. 10.1002/acp.824

Noppeney, U., Patterson, K., Tyler, L. K., Moss, H., Stamatakis, E. A., Bright, P., Mummery, C., & Price, C. J. (2007). Temporal lobe lesions and semantic impairment: A comparison of herpes simplex virus encephalitis and semantic dementia. Brain, 130(4), 1138–1147. 10.1093/brain/awl344

Noppeney, U., & Price, C. J. (2002). Retrieval of Visual, Auditory, and Abstract Semantics. NeuroImage, 15(4), 917–926. 10.1006/nimg.2001.1016

Oliver, M., Bays, R., & Zabrucky, K. (2016). False memories and the DRM paradigm: Effects of imagery, list, and test type. The Journal of General Psychology, 143, 33–48. 10.1080/00221309.2015.1110558

Oosterhof, N. N., Connolly, A. C., & Haxby, J. V. (2016). CoSMoMVPA: Multi-Modal Multivariate Pattern Analysis of Neuroimaging Data in Matlab/GNU Octave. Frontiers in Neuroinformatics, 10. https://www.frontiersin.org/articles/10.3389/fninf.2016.00027

Satterthwaite, T. D., Elliott, M. A., Gerraty, R. T., Ruparel, K., Loughead, J., Calkins, M. E., Eickhoff, S. B., Hakonarson, H., Gur, R. C., Gur, R. E., & Wolf, D. H. (2013). An improved framework for confound regression and filtering for control of motion artifact in the preprocessing of resting-state functional connectivity data. NeuroImage, 64, 240–256. 10.1016/j.neuroimage.2012.08.052

Simmonite, M., & Polk, T. A. (2022). Age-related declines in neural distinctiveness correlate across brain areas and result from both decreased reliability and increased confusability. *Aging*, Neuropsychology, and Cognition, 29(3), 483–499. 10.1080/13825585.2021.1999383

Simons, J. S., Lee, A. C. H., Graham, K. S., Verfaellie, M., Koutstaal, W., Hodges, J. R., Schacter, D. L., & Budson, A. E. (2005). Failing to Get the Gist: Reduced False Recognition of Semantic Associates in Semantic Dementia. Neuropsychology, 19(3), 353–361. 10.1037/0894-4105.19.3.353

Slotnick, S. D., & Schacter, D. L. (2004). A sensory signature that distinguishes true from false memories. Nature Neuroscience, 7(6), 664–672. 10.1038/nn1252

Slotnick, S. D., & Schacter, D. L. (2006). The nature of memory related activity in early visual areas. Neuropsychologia, 44(14), 2874–2886. 10.1016/j.neuropsychologia.2006.06.021

Spalding, K. N., Jones, S. H., Duff, M. C., Tranel, D., & Warren, D. E. (2015). Investigating the Neural Correlates of Schemas: Ventromedial Prefrontal Cortex Is Necessary for Normal Schematic Influence on Memory. The Journal of Neuroscience, 35(47), 15746–15751. 10.1523/JNEUROSCI.2767-15.2015

Stark, C. E. L., Okado, Y., & Loftus, E. F. (2010). Imaging the reconstruction of true and false memories using sensory reactivation and the misinformation paradigms. Learning & Memory, 17(10), 485–488. 10.1101/lm.1845710

Turney, I. C., & Dennis, N. A. (2017). Elucidating the neural correlates of related false memories using a systematic measure of perceptual relatedness. NeuroImage, 146, 940–950. 10.1016/j.neuroimage.2016.09.005

Tustison, N. J., Avants, B. B., Cook, P. A., Zheng, Y., Egan, A., Yushkevich, P. A., & Gee, J. C. (2010). N4ITK: Improved N3 Bias Correction. IEEE Transactions on Medical Imaging, 29(6), 1310–1320. 10.1109/TMI.2010.2046908

Vaidya, C. J., Zhao, M., Desmond, J. E., & Gabrieli, J. D. E. (2002). Evidence for cortical encoding specificity in episodic memory: Memory-induced re-activation of picture processing areas. Neuropsychologia, 40(12), 2136–2143. 10.1016/S0028-3932(02)00053-2

van Kesteren, M. T. R., Beul, S. F., Takashima, A., Henson, R. N., Ruiter, D. J., & Fernández, G. (2013). Differential roles for medial prefrontal and medial temporal cortices in schema-dependent encoding: From congruent to incongruent. Neuropsychologia, 51(12), 2352– 2359. 10.1016/j.neuropsychologia.2013.05.027

van Kesteren, M. T. R., Fernández, G., Norris, D. G., & Hermans, E. J. (2010). Persistent schema-dependent hippocampal-neocortical connectivity during memory encoding and postencoding rest in humans. Proceedings of the National Academy of Sciences, 107(16), 7550–7555. 10.1073/pnas.0914892107

van Kesteren, M. T. R., Ruiter, D. J., Fernández, G., & Henson, R. N. (2012). How schema and novelty augment memory formation. Trends in Neurosciences, 35(4), 211–219. 10.1016/j.tins.2012.02.001

Warren, D. E., Jones, S. H., Duff, M. C., & Tranel, D. (2014). False Recall Is Reduced by Damage to the Ventromedial Prefrontal Cortex: Implications for Understanding the Neural Correlates of Schematic Memory. Journal of Neuroscience, 34(22), 7677–7682. 10.1523/JNEUROSCI.0119-14.2014

Webb, C. E., & Dennis, N. A. (2019). Differentiating True and False Schematic Memories in Older Adults. The Journals of Gerontology: Series B, 74(7), 1111–1120. 10.1093/geronb/gby011

Webb, C. E., & Dennis, N. A. (2020). Memory for the usual: The influence of schemas on memory for non-schematic information in younger and older adults. Cognitive Neuropsychology, 37(1–2), 58–74. 10.1080/02643294.2019.1674798

Webb, C. E., Turney, I. C., & Dennis, N. A. (2016). What’s the gist? The influence of schemas on the neural correlates underlying true and false memories. Neuropsychologia, 93, 61–75. 10.1016/j.neuropsychologia.2016.09.023

Wheeler, M. E., Petersen, S. E., & Buckner, R. L. (2000). Memory’s echo: Vivid remembering reactivates sensory-specific cortex. Proceedings of the National Academy of Sciences, 97(20), 11125–11129. 10.1073/pnas.97.20.11125

Wise, R. J. S., & Price, C. J. (2006). Functional neuroimaging of language. In Handbook of functional neuroimaging of cognition*, 2nd ed* (pp. 191–227). Boston Review.

Yonelinas, A. P. (2002). The Nature of Recollection and Familiarity: A Review of 30 Years of Research. Journal of Memory and Language, 46(3), 441–517. 10.1006/jmla.2002.2864

Yonelinas, A. P., & Jacoby, L. L. (1995). The relation between remembering and knowing as bases for recognition: Effects of size congruency. Journal of Memory and Language, 34(5), 622–643. 10.1006/jmla.1995.1028

Zeithamova, D., Dominick, A. L., & Preston, A. R. (2012). Hippocampal and Ventral Medial Prefrontal Activation during Retrieval-Mediated Learning Supports Novel Inference. Neuron, 75(1), 168–179. 10.1016/j.neuron.2012.05.010

Zhang, Y., Brady, M., & Smith, S. (2001). Segmentation of brain MR images through a hidden Markov random field model and the expectation-maximization algorithm. IEEE Transactions on Medical Imaging, 20(1), 45–57. 10.1109/42.906424

